# Physical activity unveils the relationship between mitochondrial energetics, muscle quality and physical function in older adults

**DOI:** 10.1101/164160

**Authors:** Giovanna Distefano, Robert A. Standley, Xiaolei Zhang, Elvis A. Carnero, Fanchao Yi, Heather H. Cornnell, Paul M. Coen

## Abstract

**Background:** The concept of mitochondrial dysfunction in aging muscle is highly controversial. In addition, emerging evidence suggests that reduced muscle oxidative capacity and efficiency underlie the etiology of mobility loss in older adults. Here, we hypothesized that studying well phenotyped older cohorts across a wide range of physical activity would unveil a range of mitochondrial function in skeletal muscle and in turn allow us to more clearly examine the impact of age *per se* on mitochondrial energetics. This also enabled us to more clearly define the relationships between mitochondrial energetics and muscle lipid content with clinically relevant assessments of muscle and physical function.

**Methods:** Thirty-nine volunteers were recruited to the following study groups; Young Active (YA, n=2F/8M, age=31.2±5.4 yrs.). Older Active (OA, n=2F/8M, age=67.5±2.7 yrs.) and Older Sedentary (OS, n=8F/11M, age=70.7±4.7 yrs.). Participants completed a graded exercise test to determine fitness (VO2peak), a submaximal exercise test to determine exercise efficiency, and daily physical activity (PA) was recorded using a tri-axial armband accelerometer. Mitochondrial energetics were determined by 1) 31P magnetic resonance spectroscopy and 2) respirometry of fiber bundles from vastus lateralis biopsies. Quadriceps function was assessed by isokinetic dynamometry and physical function by the short physical performance battery (SPPB) and stair climb test.

**Results:** Daily PA energy expenditure was significantly lower in OS, compared to YA and OA groups. Despite fitness being higher in YA compared to OA and OS, mitochondrial respiration, ATPMax, P/O ratio and exercise efficiency were similar in YA and OA groups and were significantly lower in OS. P/O ratio was correlated with exercise efficiency. Time to complete the stair climb and repeated chair stand tests was significantly greater for OS. Interestingly, ATPMax was related to muscle contractile performance and physical function.

**Conclusions:** Older adults who maintain a high amount of physical activity have better mitochondrial capacity, similar to highly active younger adults, and this is related to their better muscle quality, exercise efficiency and physical performance. This suggests that mitochondria could be an important therapeutic target for sedentary aging associated conditions including sarcopenia, dynapenia and loss of physical function.

## INTRODUCTION

The progressive loss of muscle mass, strength and physical function that occurs with aging is known as sarcopenia [1, 2]. This condition is prevalent in adults over 70 years of age and predisposes an individual to mobility disability, nursing home admission and early mortality [3]. With the rapid increase in the elderly demographic in the United States, sarcopenia is poised to become a major burden on the healthcare system. Apart from resistance training, there are very few effective therapeutic strategies to treat sarcopenia [4], in part because the underlying etiology is multifactorial, complex and still needs to be completely delineated. However, there should be increased focus on developing new therapeutics for sarcopenia given its recent ICD-10-CM code designation [5].

Physical function is a key aspect of contemporary clinical definitions of sarcopenia [2], and requires the integration of multiple physiological systems, including cardiopulmonary, vestibular, sensory and muscular. A decline in skeletal muscle mitochondrial capacity also characteristic of aging and has recently been studied as an underlying factor for low physical function [6], slower walking speed [7-9], fatigability [10], and loss of exercise efficiency with aging [11]. However, it should be noted that there is still considerable debate regarding the true nature of aging-associated mitochondrial dysfunction. While many studies have supported an age-related decrease in mitochondrial capacity [12-17], several others have failed to find these associations [18-22]. These contradictory results may be explained by the fact that the majority of these studies have not considered important covariates that also affect mitochondria, such as participant physical activity (PA) levels [15], cardiovascular fitness, and adiposity [17], all of which likely confound the relationship between mitochondrial capacity and age [16, 23, 19, 24, 25]. In addition, the various analytical approaches employed to assess mitochondrial function [26], including the use of isolated mitochondria [12, 15, 17] which has been shown to exaggerate the observed deficit in mitochondrial function in aging [27], may contribute to the lack of clarity. Impaired mitochondrial function is also linked with elevated intramyocellular lipids (IMCL) in aging [28] and which have been recently linked to the pre-frail phenotype [29]. Collectively, these findings suggest that rigorously conducted investigations in well phenotyped individuals are needed to clarify the nature of age-related declines in mitochondrial function and IMCL. This is critical to further understand the relevance of mitochondrial dysfunction and IMCL in aging to clinically relevant endpoints, including those that characterize physical function.

Reduced exercise efficiency (increased oxygen consumption per work performed) for physical activities such as walking is also apparent in the elderly [30]. This in turn could impede activities of daily living in older adults [31, 32], potentially contributing to future risk for mobility disability. While the underlying cause of low exercise efficiency with aging is unknown, mitochondrial energetics may be involved [11, 33]. Furthermore, the influence of physical activity status, cardiovascular fitness and adiposity on exercise efficiency in older adults has not been thoroughly examined.

The goal of this study was to investigate the role of chronological age and PA on a comprehensive profile of skeletal muscle mitochondrial energetics, IMCL and intermuscular adipose tissue (IMAT) content. A second goal was to examine the relationship between mitochondrial energetics and IMCL with exercise efficiency, muscle quality (strength per unit mass) and physical function. The benefits of resistance exercise in preserving muscle and physical function in older adults has been relatively well documented. Here, we generated evidence from well phenotyped cohorts indicating that physical activity, predominantly endurance exercise, largely mitigates the negative effects of a sedentary lifestyle on mitochondrial energetics, exercise efficiency, and certain aspects of physical function in older adults.

## MATERIALS AND METHODS

### Study design and participants

Thirty-nine men and women from the Orlando, FL area volunteered to participate in this study. Volunteers were eligible to participate if they were weight stable (±4.5kg in preceding 6 months), had a BMI between 20-35 kg/m2 and were in good general health. Volunteers were excluded if they were taking medications known to influence muscle metabolism, had a chronic medical condition, had any contraindications to exercise, were pregnant or breast-feeding, or had high resting blood pressure (≤150mmHg systolic, ≤ 90mmHg diastolic). A note from the participant’s PCP/Cardiologist for exercise clearance was needed if positive stress test symptoms were observed from exercise test. All participants provided written informed consent and the study protocol was reviewed and approved by the institutional review board at Florida Hospital, Orlando. Thus, the study was performed in accordance with the ethical standards laid down in the 1964 Declaration of Helsinki. Participants were assigned to the following study groups; Young (21-40 yrs.) Active (YA), Older Active (OA) and Older Sedentary (OS, 65-90 yrs.). Participants were considered active if they engaged in aerobic exercise (running, cycling, swimming) at least 3 days a week without extensive layoff over the previous 6 months. Participants were considered sedentary if they completed one or less structured exercise sessions a week.

All testing was completed over four study visits at the Translational Research Institute for Metabolism and Diabetes at Florida Hospital, Orlando. Visit #1 consisted of a fasting blood draw, physical measurements, medical history/physical activity questionnaires, resting ECG. On visit #2 participants completed a VO2 peak test with ECG to determine cardiorespiratory fitness. On visit #3, participants completed magnetic resonance imaging (MRI)/ magnetic resonance spectroscopy (MRS), quadriceps contractile and physical function testing, dual energy x-ray absorptiometry (DXA) scan and were given a triaxial accelerometer to determine free living physical activity over a 1-week period. On study visit #4, fasting blood samples were drawn and resting energy expenditure was determined by indirect calorimetry. Participants then consumed a small low glycemic index meal (200 kcal, 15% protein, 35% fat, 50% carbohydrate) and 15 minutes later the muscle biopsy procedure was conducted.Participants then completed the acute exercise bout on a cycle ergometer, all as described below.

### Daily physical activity

The monitor used for this study was the SenseWear® Pro Armband (BodyMedia Inc., Pittsburgh, PA). This activity monitor uses 3-axis accelerometer, galvanic skin response and skin temperature to estimate energy expenditure at a one-minute resolution [34]. The SenseWear® Armband has been demonstrated to be a valid, accurate and reliable method when compared to indirect calorimetry [34], and doubly labeled water [35]. Participants were instructed to wear activity monitors on their upper left arm for at least seven consecutive days. Only days with a wear time of at least 85% were considered for further analysis.

### Cardiorespiratory fitness

VO2 peak was determined as peak aerobic capacity measured during a graded exercise protocol on an electronically braked cycle ergometer, as previously described [7, 36]. Heart rate, blood pressure and ECG (12-lead) was recorded throughout this test. The test was terminated according to the criteria outlined in the American college of sports medicine exercise testing guidelines [37].

### Assessment of body composition

<u>Dual energy X-ray absorptiometry (DXA)</u> scans were performed using a GE Lunar iDXA whole-body scanner and analyzed with enCORE software. Coefficients of variation (CV) vary by tissue type and range from 0.5% to 1.1%. <u>Magnetic resonance imaging</u> (MRI) was performed on a 3T Philips Acheiva magnet and was used to assess of intermuscular (IMAT) and subcutaneous fat (SAT). Volumetric measurement of fat was performed under standardized conditions with subjects in a supine position. T1 weighted imaging sequences were performed across the thighs. The resultant images were analyzed using Analyze 11.0 (Biomedical Imaging Resource, Mayo Clinic, Rochester MN) to segment IMAT and SAT depots and to measure the muscle volume. CVs based on same day measurements with repositioning between scans are 1.1%, 4.2% and 0.4% for SAT, IMAT and muscle, respectively.

### Magnetic resonance spectroscopy (MRS)

All MRS measurements were acquired on a 3T Philips Acheiva magnet. <u>Assessment of Intramyocellular lipids</u> (IMCL) was by proton (^1^H) spectroscopy using multiple single voxel acquisitions. Ratios of IMCL to water were determined using the AMARES (Advances Method for Accurate, Robust and Efficient Spectral fitting) algorithm within the jMRUI software. CVs based on same day measurements with repositioning between scans are 7.6% in the soleus and 18.3% in the tibialis anterior. <u>Assessment of ATP</u> _Max_ was performed as previously described [38, 39].The Phosphocreatine (PCr) recovery time constant (tau) and the PCr level in oxygenated muscle at rest (PCr) _rest_ were used to calculate maximum mitochondrial capacity (ATP _Max_). After the fully relaxed spectrum (TR 15s, TE 0.17s, NSA 48) was obtained, the ATP_Max_ experiment was performed by obtaining 31P spectra, with one summed spectrum acquired every 6 seconds for the duration of the ATP_Max_ experiment. Briefly, a ^31^P surface coil was used to measure phosphorous containing species including PCr, ATP and Inorganic phosphate (P_i_) using a standard one pulse acquisition experiment. For the first minute, the participant remained still to obtain baseline measurements. After the baseline was established, the volunteer was asked to perform isometric contractions of the quadriceps (by slight kicking) for up to 45 seconds. After kicking stopped, the volunteer remained still for an additional ~8 minutes to allow the PCr peak to return to baseline. The entire ATP_Max_ acquisition took 10 minutes. The experimental spectra (6sec each) consisted of 4 (NSA) partially saturated free induction decays (TR = 1.5s, TE = 0.13s). The areas/heights of the peaks in each ^31^P MRS spectrum during the experiment were determined using the AMARES (Advances Method for Accurate, Robust and Efficient Spectral fitting) algorithm within the jMRUI software. Established formulas were used to determine the intracellular pH based on the PCr and Pi chemical shifts and to calculate tau [40]. The CV for the ATP_Max_measured in the same participant in one day repositioned or with less than a week between measurements was on average 3.6%.

### Muscle contractile performance

Quadriceps contractile performance of the left thigh was determined during a leg extension exercise using an isokinetic pneumatic-driven dynamometer equipped with load cells and potentiometers (Biodex Medical Systems, Inc., Shirley, NY). One repetition maximum for left leg knee extension was determined using a standard weight stack. The initial weight was determined as a predicted 1 repetition maximum (1-RM), where the starting weight was 90% and 60% of body weight for males and females, respectively. For each additional 10 years of age the 1-RM decreased 10%.

### Physical function testing

Lower extremity function was assessed by short physical performance battery (SPPB), which consisted of three tasks: five repeated timed chair stands, timed standing balance (with feet in parallel, semi-tandem, and tandem positions), and a 4-meter walk to determine usual gait speed [41]. Additionally, each participant completed a timed stair climb test and a grip strength test using a Jamar® hand-held digital dynamometer.

### Percutaneous skeletal muscle biopsy

Percutaneous muscle biopsies were obtained in the morning and participants were instructed not to perform physical exercise 48 hours prior to the biopsy procedure. Prior to the biopsy resting metabolic rate was determined while resting in the supine position over 30 mins by indirect calorimetry using a canopy system (Parvo Medics, Sandy, UT). Biopsy samples were obtained from the middle region of the vastus lateralis under local anesthesia (2% buffered lidocaine) as described previously [42]. A portion of the biopsy specimen was placed in ice-cold preservation buffer (BIOPS) for analysis of mitochondrial respiration [7].

### Submaximal exercise test

Participants performed a 6-minute warm-up consisting of light cycling on an electronically braked cycle ergometer. After the warm-up, subjects cycled for 40 minutes at approximately 70% of heart rate reserve (HRR) which was calculated as HRmax during the VO_2_peak test minus supine resting HR. Heart rate, perceived exertion and blood pressure data were collected every 5 mins during the test. Indirect calorimetry measurements were recorded using an open circuit spirometry metabolic monitor system (Parvo Medics, Sandy, UT). Respiratory variables were collected to verify that the participant was exercising at the correct intensity, which was achieved within 5 minutes of starting the test, and to calculate exercise efficiency parameters.

### High-resolution respirometry and ROS emission

Permeabilized myofiber bundles (~1-3mg each) were prepared immediately after the biopsy procedure, as previously described [43]. Briefly, individual fibers were gently teased apart in a petri dish containing ice-cold BIOPS media. The fiber bundles were then permeabilized with saponin (2ml of 50μg/ml) for 20 minutes and washed twice (10 minutes each) in Buffer Z (105mM K-MES, 30mM KCl, 10mM KH_2_PO4, 5mMMgCl_2_-6H_2_O, 5mg/ml BSA, 1mM EGTA,pH7.4 with KOH). Mitochondrial respiration was evaluated by high-resolution respirometry (Oxygraph-2k, Oroboros Instruments, Innsbruck, Austria). Measurements were performed at 37°C, in the range of 400-200 nmol O_2_/ml. Two assay protocols were run in duplicate using Buffer Z with blebbistatin (25μM). In protocol 1, complex I supported LEAK (L_I_) respiration was determined through the addition of malate (2mM) and glutamate (5mM). ADP (4mM) was added to elicit complex I supported oxidative phosphorylation (OXPHOS) (P_I_) respiration. Succinate (10mM) was then added to elicit complex I and II supported OXPHOS (P_I+II_) respiration. Cytochrome c (10μM) was added to assess the integrity of the outer mitochondrial membrane. Finally, a titer of FCCP (carbonyl cyanide-p-trifluoromethoxyphenylhydrazone) (2μM steps) was performed to determine the maximal electron transfer system (E_I+II_) capacity or maximal uncoupled respiration. In protocol 2, FAO supported LEAK (FAO_L_) respiration was determined through the addition of palmitoylcarnitine (25μM) and malate (2mM). ADP (4mM) was added to elicit FAO supported OXPHOS (FAO_*P*_). Cytochrome c (10μM) was added to access mitochondrial integrity. Glutamate (5mM) was then added to elicit complex I and FAO_*P*_ (CI&FAO_*P*_) respiration. Succinate was added (10mM) to stimulate complex I and II and FAO_*P*_ (CI+II&FAO_*P*_). Steady state O_2_ flux for each respiratory state was determined and normalized to myofiber bundle wet weight using Datlab 5 software (Oroboros Instruments, Innsbruck, Austria).

Measurement of H_2_O_2_ emission by mitochondria was measured in from permeabilized muscle fiber bundles by real time monitoring of Amplex Red oxidation using a SPEX Fluoromax 3 (HORIBA Jobin Yvon) spectrofluorometer with temperature control and magnetic stirring as previously described [44]. Mitochondrial H_2_O_2_ emission rate was expressed as pmol/min/mg wet weight.

### Exercise efficiency calculations

Exercise efficiency was calculated from calorimetry data generated during the submaximal test, as previously described [45]. Energy expenditure during steady-state exercise (20 to 40 minutes) was calculated using the Brouwer equation as previously described [46]. Gross efficiency (GE) was calculated as the ratio of work output to exercise energy expenditure during the submaximal exercise bout, as follows: GE (%) = (Work _(kcal/min)_/EE_(kcal/min)_)^*^100. Net Efficiency (NE) was determined as the ratio of work output to exercise minus resting energy expenditure, as follows: NE (%) = (Work_(kcal/min)_/EE_(kcal/min)_-REE_(kcal/min)_)^*^100.

### Statistics

Group differences were determined using a one-way ANOVA with a Tukey’s post hoc test for individual group comparisons. The distribution of sex and race across groups were determined by Chi-squared test. To measure associations between variables while adjusting for age and gender, the partial correlation analyses were conducted. The forward stepwise and multiple regression models were conducted to explore the best combination of variables to explain muscle and physical function. Statistical significance was set at P<0.05. Statistical analyses were performed using SAS 9.4 and JMP v9.0 (SAS, Cary, NC).

## RESULTS

### Participants characteristics

Thirty-nine participants were screened and enrolled to one of three groups: Young Active (YA, n=10), Old Active (OA, n=10), or Old Sedentary (OS, n=19). The characteristics of the study groups are shown in table 1. By study design, the OS group had lower objectively measured daily energy expenditure (EE), activity EE and steps per day, compared to the active groups. The OS group had lower cardiovascular fitness and greater BMI compared to the active groups indicating deconditioning and increased adiposity typically associated with a sedentary lifestyle. Weight and sex ratio were similar between groups. The results of the DXA and MRI scans and representative mid-thigh MR images are presented in table 1 and figure 1, respectively. Whole body fat mass, and mid-thigh SAT and IMAT volumes were greater in the OS group, indicating greater adiposity. While whole body lean mass was similar between groups (P = 0.065, table 1), left mid-thigh lean mass (table 1) and thigh muscle volume were significantly lower in the OS group when compared to the YA group (figure 1). Skeletal muscle index (SMI) was similar between groups and above sex-specific population based cutoffs indicating the absence of low muscle mass and thus sarcopenia [2]. Hand grip strength was lower in both older groups, compared to the YA group.

**Table 1.** Participant fitness, physical activity and body composition characteristics.

|  | **YA (n=10)** | **OA (n=10)** | **OS (n=19)** | **ANOVA**<br>**P-Value** |
| --- | --- | --- | --- | --- |
| **Age, years** | 31.2±5.4 <sup>B</sup> | 67.5±2.7 <sup>A</sup> | 70.7±4.7 <sup>A</sup> | <0.0001 |
| **Sex, Ratio** | 2F/8M | 2F/8M | 8F/11M | 0.327* |
| **Weight, Kg** | 67.7±9.7 | 69.9±15.1 | 79.1±3.1 | 0.075 |
| **BMI, Kg/m<sup>2</sup>** | 21.3±1.0 <sup>B</sup> | 24.2±1.0 <sup>B</sup> | 27.7±0.7 <sup>A</sup> | <0.0001 |
| **Waist, cm** | 74.6±5.1 <sup>B</sup> | 88.2±12.3 <sup>A</sup> | 97.4±11.9 <sup>A</sup> | <0.0001 |
| **VO<sub>2</sub> peak (ml/kgBW/min)** | 56.7±9.6 <sup>A</sup> | 34.9±4.3 <sup>B</sup> | 18.58±3.8 <sup>C</sup> | <0.0001 |
| **VO<sub>2</sub> peak (ml/kgFFM/min)** | 65.8±3.4 <sup>A</sup> | 45.3±1.8 <sup>B</sup> | 29.6±0.8 <sup>C</sup> | <0.0001 |
| **Daily EE (kcal/24hr)** | 3108± 270 <sup>A</sup> | 2565±555 <sup>B</sup> | 2117±404 <sup>C</sup> | <0.0001 |
| **Daily AEE (kcal/24hr)** | 1833±369 <sup>A</sup> | 1248±545 <sup>B</sup> | 488±335 <sup>C</sup> | <0.0001 |
| **Steps per day (steps/24hr)** | 10560±4242 <sup>A</sup> | 8459±2991 <sup>A</sup> | 4883±2683 <sup>B</sup> | 0.0003 |
| **PASE Score** | - | 215±25 | 105±18 | 0.0017 |
| **Total Fat Mass, Kg** | 10.1±0.9 <sup>B</sup> | 17.3±2.8 <sup>B</sup> | 29.9±1.7 <sup>A</sup> | <0.0001 |
| **Total Lean Mass, Kg** | 55.4±2.6 | 51.5±2.9 | 46.6±2.3 | 0.065 |
| **Left Leg Lean Mass (Kg)** | 9.5±0.50 <sup>A</sup> | 8.3±0.46 <sup>AB</sup> | 7.7±0.43 <sup>B</sup> | 0.033 |
| **SMI (ALM/h<sup>2</sup>)** | 8.23±0.3 | 7.89±0.3 | 7.43±6.9 | 0.1859 |
| **Hand Grip Strength (Kg)** | 44.3±2.8 <sup>A</sup> | 35.6±2.5 <sup>B</sup> | 31.9±1.7 <sup>B</sup> | 0.001 |

Data are presented as Mean±SD. <sup>A, B and C</sup> indicate significant differences between groups (p<0.05). \* Sex distribution across groups were determined by Chi Square test. Abbreviations: BMI, Body mass index; BW, Body weight; FFM, Fat-free mass; EE, Energy expenditure; AEE, Activity energy expenditure; PASE, Physical activity scale for the elderly; SMI, Skeletal muscle index; ALM, Appendicular lean mass.

**Figure 1.**
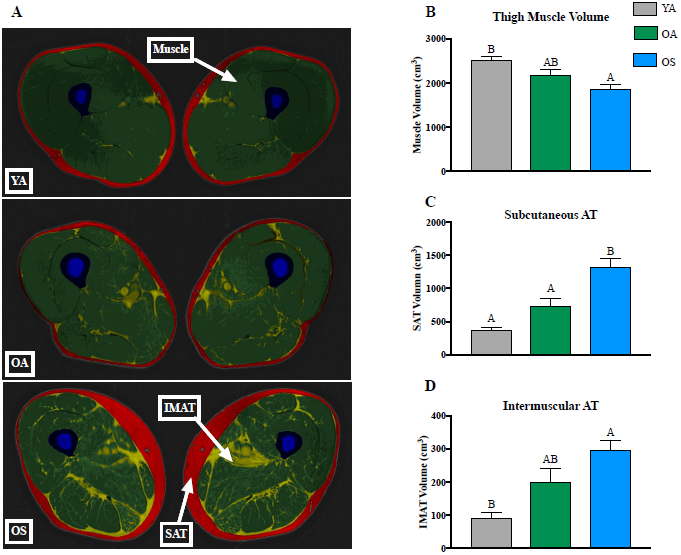
Mid-thigh muscle and adipose tissue volume. <u>Panel A</u>, Representative mid-thigh MR imagesfrom young active (YA), old active (OA), and old sedentary (OS) participants. Skeletal muscle is colored dark green. Subcutaneous adipose tissue (SAT) is colored red. Intermuscular adipose tissue (IMAT) is colored light green. <u>Panel B</u>, Mid-thigh muscle volume was higher in the YA group when compared to OS. Physical activity partially preserved mid-thigh muscle volume in the OA group. <u>Panel C</u>,Subcutaneous adipose tissue (SAT) volume was higher in the OS group when compared to YA and OA groups. <u>Panel D</u>, Intermuscular adipose tissue (IMAT) volume was higher in the OS group when compared to YA. Physical activity partially prevented IMAT accumulation in the OA group. The letters A and B denote significant differences between groups (P < 0.05, One-way ANOVA followed by post-hoc Tukey test). Data presented are Mean ± SEM.

### Muscle contractile performance and physical function

Left leg peak torque, total work, average power and 1-repitition maximum were greater in the YA group compared to both older groups (table 2 and figure 2A). To determine if differences in leg skeletal muscle mass between groups may explain differences in muscle contractile performance, leg 1-RM was normalized to left leg lean mass (from regional DXA), to produce an index of muscle quality (figure 2B). The YA group had significantly greater normalized 1-RM compared to the OS group. However, OA had similar normalized 1-RM to YA indicating that PA rescued some aspects of age-associated decline of muscle quality.

**Table 2.** Muscle Contractile performance by isokinetic dynamometry.

|  | **YA (n=10)** | **OA (n=10)** | **OS (n=19)** | **ANOVA**<br>**P-Value** |
| --- | --- | --- | --- | --- |
| **BIODEX (Left Leg, 120°/sec)** |  |  |  |  |
| **Peak Torque (Nm)** | 114±11.2 <sup>A</sup> | 99±8.4 <sup>B</sup> | 87±5.6 <sup>B</sup> | <0.0001 |
| **Total Work (W)** | 929±79 <sup>A</sup> | 594.4±50 <sup>B</sup> | 532±38 <sup>B</sup> | <0.0001 |
| **Average Power (J)** | 195±16 <sup>A</sup> | 128±10 <sup>B</sup> | 111±8.6 <sup>B</sup> | <0.0001 |
| **1 Repetition Maximum (Kg)** | 51±4.6 <sup>A</sup> | 36±3.4 <sup>B</sup> | 26±2.4 <sup>B</sup> | <0.0001 |

Data: Mean±SD for all subjects. <sup>A, B</sup> indicate significant differences between group (p<0.05).
Abbreviations: Nm, newton meters; W, Watt; J, joule; sec, second.

**Figure 2.**
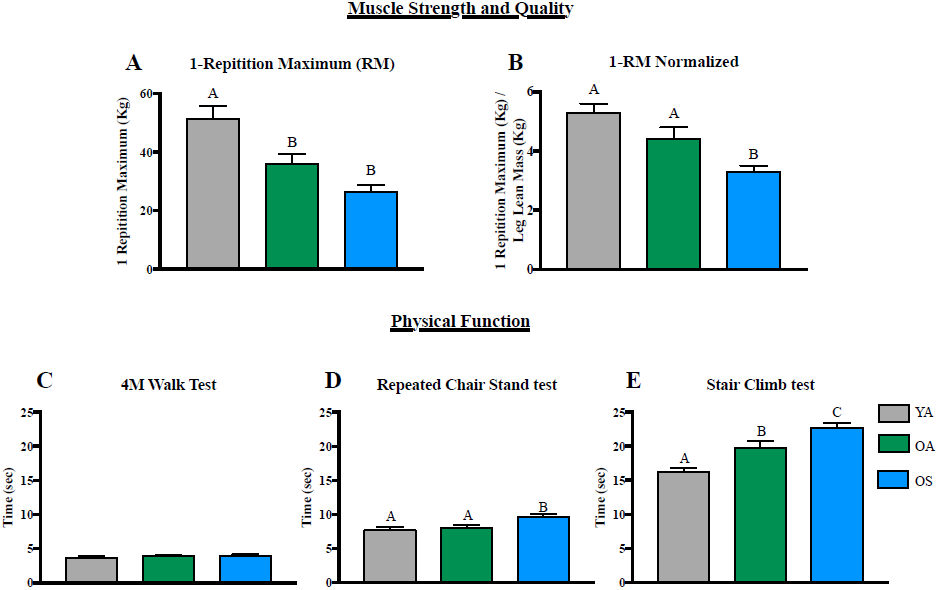
Muscle contractile performance and physical function. <u>Panel A</u>, Left leg 1-repititionmaximum (1-RM) was lower in the OA and OS groups, compared to YA. <u>Panel B</u>, Muscle quality (1-RM/left leg lean mass) was similar between YA and OA groups, and lower in the OS group. Physical function was assessed *via* 4m walk test (from SPPB), repeated chair stand (from SPPB), and stair climb test. <u>Panel C</u>, There was no difference between groups in time to complete 4m walk test. <u>Panel D</u>, Time to complete the repeated chair stand test was higher in the OS group when compared to the YA and OA groups. <u>Panel E</u>, Time to complete the stair climb test was higher in the OS group when compared to YA and OA groups, and higher in the OA when compared to YA. The letters A, B and C denote significant differences between groups (P < 0.05, One-way ANOVA followed by post-hoc Tukey test). Data presented are Mean ± SEM.

There was no difference in the SPPB score between OA and OS (11.9 vs. 11.5, P >0.05).However, when the individual test results were considered, group differences were apparent. The results are presented in figure 2. Interestingly, the duration of the test seemed to differentiate physical function between the three groups. There was no difference between groups for the time taken to walk 4m (figure2C). Repeated chair stand took longer to complete for the OS group, when compared to the active groups (figure 2D). Finally, time to complete the stair climb test was significantly different between all three groups, with the OS group having the poorest performance (figure 2E).

### Skeletal muscle mitochondrial energetics are affected by physical activity and not chronological age

We employed ^31^P MRS to examine in-vivo mitochondrial energetics. This approach assesses maximal capacity of the mitochondrial to generate ATP (ATPmax) following muscle contraction and integrates all aspects of mitochondrial content and function including oxygen consumption, Ca^2+^ handling and redox, all working together to produce ATP [47]. We observed that both YA and OA groups had similar maximal ATP synthetic rate and PCr recovery time following muscle contraction and that the OS group had poorer mitochondrial energetics (figure 2A&B). These results indicate that maximal ATP synthesis of muscle is impacted by physical activity (or inactivity) to a greater degree than chronological age *per se*.

To complement ^31^P MRS measurements, we employed an ex-vivo approach to assess respiratory capacity of the electron transfer system (ETS) in muscle biopsy specimens. Measuring respiration in permeabilized muscle fibers is a widely-employed approach to specifically assess mitochondrial ETS function. In the first assay protocol, which utilized carbohydrate-derived substrates, we found that OS had lower LEAK (L_I_), CI and CI+II supported OXPHOS respiration (P_I_ and P_I+II_, respectively), and maximal ETS capacity (E_I+II_) compared to the OA and YA groups which were similar (figure 3C). The second assay protocol used fatty acid (palmitoyl-carnitine) and carbohydrate-derived substrates. Again, we observed that the OS group had lower LEAK (L_FAO_), and CI+II+FAO supported OXPHOS (P _I+II+FAO_) respiration (figure 3D). Finally, the cytochrome C response for all groups was <10%, indicating maintenance of mitochondrial membrane integrity. These finding support the observation of lower ATP_Max_ in the OS group and that is likely due to reduced ETS respiratory capacity. These finding clearly indicate that the etiology of lower mitochondria energetics in aging muscle primarily due to low physical activity.

**Figure 3.**
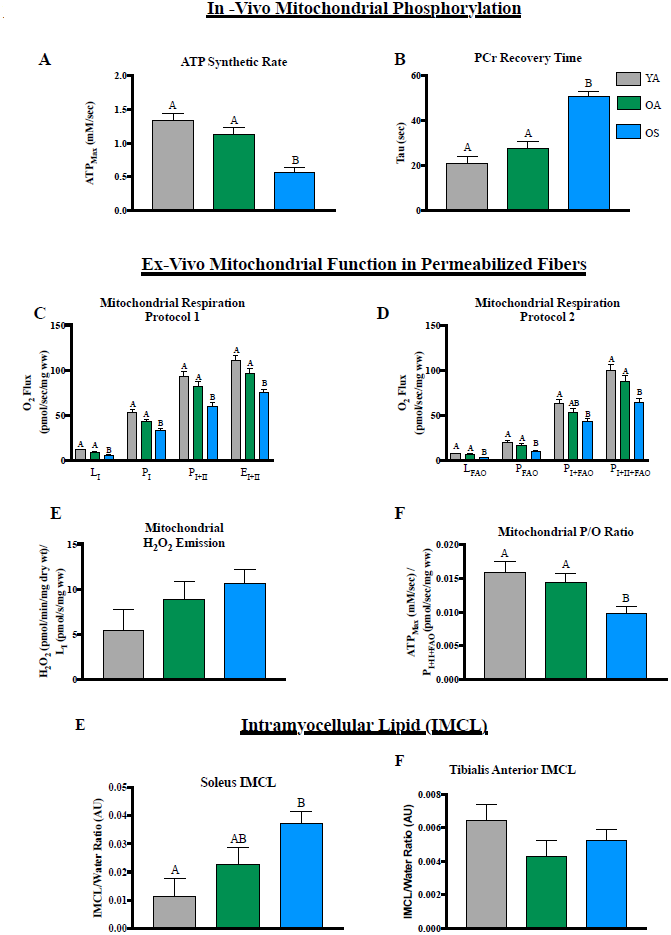
Skeletal muscle mitochondrial energetics and intramyocellular lipid. Maximal ATP synthetic rate (ATP_Max_) and phosphocreatine (PCr) recovery time were determined by 31P MRS. <u>Panel A</u>,ATP_Max_ was lower in OS compared to both YA and OA groups. <u>Panel B</u>, PCr recovery time was significantly longer for OS compared to OA and YA groups. Mitochondrial respiratory capacity measured in permeabilized myofibers by high-resolution respirometry. <u>Panel C</u>, OS had lower LEAK respiration (L_I_, 5mM glutamate & 2mM Malate), ADP (4mM) stimulated maximal complex I (P_I_) and complex I and II (P_I+II_, 10mM succinate) OXPHOS respiration, and maximal electron transfer system capacity (EI+II, FCCP titer 0-2μM), when compared to YA and OA groups. <u>Panel D</u>, OS had lower fatty acid supported LEAK respiration (LFAO, 2mM Malate & 25uM Palmitoylcarnitine), ADP (4mM) stimulated maximal OXPHOS respiration supported by FAO (PFAO,), and complex I and II supported FAO (P_I+II+FAO_, 2mM Malate, 5mM Glutamate & 10mM succinate), when compared to OA and YA groups. <u>Panel E</u>,Mitochondrial H_2_O_2_ emission in permeabilized myofibers was not different between groups. <u>Panel F</u>,Mitochondrial efficiency (ATP_Max_ / P_I+II_respiration; P/O ratio) was lower in the OS group when compared to YA and OA groups. <u>Panels G and H</u>, Skeletal muscle intramyocellular lipid (IMCL) content.<u>Panel G</u>, Soleus IMCL accumulation was higher in the OS when compared to YA. Physical activity partially prevented soleus IMCL accumulation in the OA group. <u>Panel F</u>, Tibialis anterior IMCL content was similar between groups. The letters A&B denote significant differences between groups (P < 0.05, One-way ANOVA and Tukey’s honest significance difference post-hoc test). Data presented are Mean ± SEM.

Mitochondria are also a source of redox signaling, an aspect of mitochondrial function that is reported to be dysfunctional with high fat feeding, obesity and aging. We assessed mitochondrial H_2_O_2_ emission in permeabilized fiber bundles via real time fluorometric monitoring of the Amplex Red reaction. We found that ROS emission tended to be higher in the OS group, but did not reach statistical significance (figure 3E). We also examined whether an index of mitochondrial efficiency (P/O; ATP_Max_ – maximal ATP synthesis / PI+II – maximal mitochondrial oxygen consumption) was different between groups. We previously reported that this index of mitochondrial efficiency was related to 400m walk time in older adults [7]. Here we show that older sedentary individuals have lower mitochondrial P/O ratio compared to both active groups (figure 3F).

### Soleus intramyocellular lipid is elevated in older sedentary individuals

Mitochondrial dysfunction in aging often occurs in parallel with accumulation of intramyocellular lipid, which in turn may contribute to dysfunctional muscle metabolism. We used proton MR spectroscopy to assess intramyocellular lipid in soleus and tibialis anterior muscle groups (figure 3G&H). Interestingly, IMCL was elevated in the soleus of the OS group compared to YA, while tibialis anterior IMCL content was not different between groups. This result raises the interesting question that age and physical activity may have muscle group specific effects on IMCL, and potentially mitochondrial function, which in turn may be linked to the distinct muscle fiber compositions of these muscle groups.

### Mitochondrial and exercise efficiency are related and both are reduced with physical inactivity and not chronological age

A reduction in work (or exercise) efficiency has been reported to contribute to increased cost of movement (walking, cycling) in older adults. We measured exercise efficiency during a submaximal exercise bout to examine the impact of physical activity and aging. The group average HRR that was achieved during the exercise bout was 72% (YA), 71.5% (OA) and 68% (OS) percent and this equated to 61% (YA), 63% (OA), and 64% (OS) of VO2 peak (ml/Kg/min) indicating all three groups exercised at the same relative workload. We found that both gross (GE) and net (NE) efficiencies were similar between the active groups and lower in the OS group (figure 4A&B). Interestingly, when we examined data from all subjects (n=39) we found that mitochondrial P/O ratio was also positively related to both GE (R^2^ = 0.22, P = 0.01) and NE (R^2^ = 0.13, P = 0.049), indicating a link between mitochondrial efficiency in muscle and whole body exercise efficiency. In addition, PCr recovery time following contraction was negatively related to GE (R^2^ = 0.22, P = 0.01) and NE (R^2^ = 0.13, P = 0.049), suggesting that mitochondrial ATP synthetic capacity is related to exercise efficiency.

**Figure 4.**
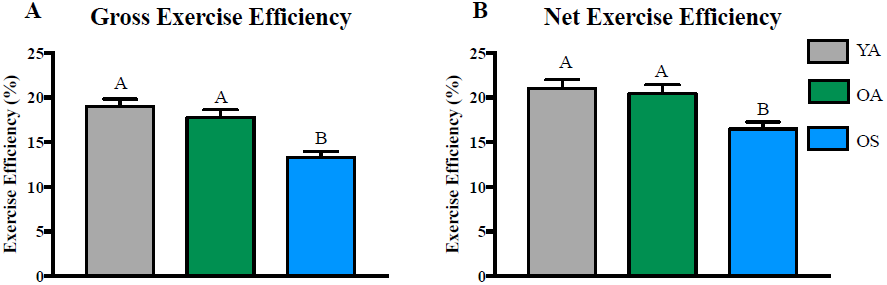
Exercise efficiency and its association with mitochondrial energetics and intermuscular adipose tissue. The OS group had lower Gross (<u>Panel A</u>) and Net (<u>Panel B</u>) exercise efficiency compared to the active groups. The letters ^A&B^ denote significant differences between groups (P < 0.05, One-way ANOVA and Tukey’s honest significance difference post-hoc test). Data presented are Mean ± SEM.

### Mitochondrial energetics and IMCL relate to muscle quality and physical function in older adults

To further explore the relationship between mitochondrial energetics, muscle quality and physical function, we conducted partial correlations while controlling for the effects of age and sex in the OA and OS groups only (table 3, n = 29). We found that muscle quality (1RM/leg lean mass) was independently explained by mitochondrial energetics (ATP_Max_, Tau, respiration, P/O ratio) and muscle lipid content (IMAT, soleus IMCL) variables. Interestingly, soleus IMCL content was also associated with gait speed, step test time and repeated chair stand time.

**Table 3.** Partial correlations.

|  |  | **1 RM**<br>**((Kg)/(kg))** |  | **Gait speed**<br>**(m/sec)** |  | **Step Test (sec)** |  | **Repeated Chair Stand**<br>**(sec)** |  |
| --- | --- | --- | --- | --- | --- | --- | --- | --- | --- |
| **Measurement** | **Variables** | **r** | **p** | **r** | **p** | **r** | **p** | **r** | **p** |
| **Mitochondrial energetics** | **ATP<sub>Max</sub> (mM/sec)** | 0.457 | **0.016\*** | 0.303 | 0.133 | -0.046 | 0.818 | -0.188 | 0.358 |
|  | **Tau (sec)** | -0.334 | **0.088** | -0.236 | 0.246 | 0.131 | 0.516 | 0.210 | 0.304 |
|  | **Maximal coupled respiration**<br>**(pmol/sec/mg ww)** | -0.016 | 0.939 | -0.182 | 0.395 | -0.192 | 0.358 | -0.054 | 0.803 |
|  | **H<sub>2</sub>O<sub>2</sub> (pmol/min/mg dw)** | 0.389 | **0.054\*** | -0.076 | 0.724 | -0.329 | 0.108 | -0.271 | 0.200 |
|  | **P/O ratio**<br>**((mM/sec)/(pmol/sec/mg ww))** | 0.545 | **0.007\*** | 0.263 | 0.226 | 0.276 | 0.202 | 0.114 | 0.606 |
| **Muscle lipid content** | **IMAT volume (cm<sup>3</sup>)** | -0.487 | **0.010\*** | -0.372 | **0.061** | 0.253 | 0.203 | 0.348 | **0.081** |
|  | **Soleus IMCL (AU)** | -0.530 | **0.004\*** | -0.594 | **0.001\*** | 0.345 | **0.078** | 0.388 | **0.050\*** |
| **Cardiovascular fitness and Physical activity** | **VO<sub>2</sub> peak (L/min)** | 0.165 | 0.410 | 0.235 | 0.248 | -0.338 | **0.085** | -0.362 | **0.069** |
|  | **Steps per day (steps/24h)** | 0.294 | 0.154 | 0.398 | **0.054\*** | -0.351 | **0.085** | -0.301 | 0.153 |
| **Exercise efficiency** | **Gross efficiency (%)** | 0.411 | **0.046\*** | 0.244 | 0.251 | -0.239 | 0.260 | -0.322 | 0.126 |
|  | **Net efficiency (%)** | 0.340 | 0.104 | 0.188 | 0.379 | -0.211 | 0.323 | -0.244 | 0.251 |

Data for partial correlations are from the OA and OS groups only (total n = 29). Correlations are adjusted for Age and Sex. \* indicate statistically significant partial correlation (p<0.05) and bold p-values indicate p<0.10. Abbreviations: ATP<sub>Max</sub>, maximal mitochondrial phosphorylation capacity; IMAT, intermuscular adipose tissue; IMCL, intramyocellular lipid; RM, repetition maximum.

We next constructed multiple regression models to more deeply explore how combinations of variables explain variation in muscle quality and physical function (table 4). While including age and sex as covariates, we found that ATP_Max_ and soleus IMCL explained 60.6% of the variation in muscle quality. Soleus IMCL was the single strongest predictor of gait speed explaining 30.3% of variation. Finally, 54.6% of the variation in time to complete step test was explained by a combination of ATP_Max_, leg lean mass and VO_2_peak.

**Table 4.** Regression models.

| **Parameter** | **1 RM (kg/kg)** |  | **Gait speed (m/sec)** |  | **Step test (sec)** |  |
| --- | --- | --- | --- | --- | --- | --- |
| **Adj $R^2$ (Model $p$)** | 0.606 (<0.0001) |  | 0.303 (0.0083) |  | 0.546 (0.0002) |  |
| | $\beta$ | p | $\beta$ | p | $\beta$ | p |
| **Age (yrs)** | -0.336 | 0.017* | 0.09 | 0.596 | -0.007 | 0.963 |
| **Sex** | -0.465 | 0.004* | 0.258 | 0.153 | -0.037 | 0.864 |
| **ATP<sub>Max</sub> (mM/sec)** | 0.319 | 0.03* | - | - | 0.656 | 0.003 |
| **Soleus IMCL (AU)** | -0.349 | 0.008* | -0.581 | 0.001* | - | - |
| **Leg lean mass (Kg)** | - | - | - | - | 1.276 | <0.001 |
| **VO<sub>2</sub>peak (L/min)** | - | - | - | - | -1.746 | <0.001 |

Data for multiple regression models are from the OA and OS groups only (total n = 29). \* indicate statistically significant contribution to the multiple regression model ( $p < 0.05$ ). Abbreviations: ATP<sub>Max</sub>, maximal mitochondrial phosphorylation capacity; IMCL, intramyocellular lipid; RM, repetition maximum.

## DISCUSSION

The paradigm of age-associated mitochondrial dysfunction in human skeletal muscle continues to generate considerable debate. In addition, the clinical relevance of mitochondrial dysfunction in aging has not been clearly defined. A deeper understanding is critical if mitochondria are to be a feasible therapeutic target for age-related conditions including sarcopenia and loss of physical function. We rationalized that studying well phenotyped older cohorts across a wide range of physical activity, principally endurance exercise, and comparing to a young active group, would unveil a range of mitochondrial function in skeletal muscle. This in turn allowed us to more clearly examine the impact of age *per se* on mitochondrial energetics and the relationships between mitochondrial energetics with clinically relevant assessments of physical function. Indeed, the OS group steps/day average was below the 5000 step threshold for sedentary behavior for healthy adults [48]. Here, we show that PA largely mitigates the negative effects of a sedentary lifestyle on mitochondrial energetics, exercise efficiency and certain aspects of muscle and physical function in older adults. Furthermore, novel evidence obtained at the myocellular and whole body level indicate that mitochondrial energetics is a determinant of exercise efficiency, muscle quality and physical function.

### The influence of chronological age and physical activity on physical function and muscle quality

Age-associated loss in the capacity for mobility is an increasingly important focus in the field of gerontology. While the consensus is that the etiology of mobility loss is multifactorial, much work remains to identify specific key physiological factors that underlie mobility loss with aging. Among the physical function tests used, we found that the longer duration of the stair climb test effectively separated the three groups, whereas the short duration 4m walk test did not. These results are in line with the observations of Choi et al., and suggest that maintaining PA into old age, and as a result cardiovascular fitness and muscle bioenergetics, preserves physical function in longer more difficult tests of physical function [8]. Interestingly, age-associated loss of muscle strength (1-RM) and muscle contractile performance assessed by isokinetic dynamometry was evident, despite a physically active lifestyle in the OA group. However, in our study PA did seem to partially preserve muscle quality (strength per muscle mass). The lack of preservation of strength with endurance type PA may be due to a relatively lower engagement of type II fibers during endurance exercise (running, swimming, cycling) compared to type I fibers, that in turn may not be sufficient to protect against age-associated preferential atrophy of type II fibers [49]. In this respect, our data support the notion that resistance exercise may be needed to effectively offset loss of muscle strength with aging.

### Physical Activity, not chronological age, influences mitochondrial energetics of muscle

A key finding was that mitochondrial energetics of skeletal muscle was greatly influenced by physical activity status and not chronological age. There are numerous reports of decrements in mitochondrial oxidative capacity with aging [12-17]. However, it is important to note that many of these investigations do not characterize important aspects of participant phenotype that likely influence the relationship between age and mitochondrial function, including objectively measured daily physical activity level and cardiorespiratory fitness [15, 14]. Many have also used isolated mitochondria [12, 15, 17], an experimental approach that may confound assays of mitochondrial function in aging [50].

In contrast to the aforementioned studies, others who control for participant phenotype have failed to find age-related changes in mitochondrial respiration in permeabilized myofibers [19, 20, 22, 51, 18]. Our study is the first to comprehensively interrogate mitochondrial energetics (in vivo and ex vivo) in active young and older groups who are well phenotyped in terms of objectively measured physical activity, cardiovascular fitness, and body composition. Our results are in line with two recent reports showing that physically active young and old groups have similar protein markers of mitochondrial content and respiration, data that underscore the importance of considering physical activity when studying mitochondrial function in aging [18, 52]. Our findings also extend this work and suggest that lower ATP synthesis capacity in the OS group is likely due to reduced ETS respiratory capacity. We also assessed mitochondrial H_2_O_2_ emission in permeabilized fiber bundles and found that ROS emission tended to be higher in the OS group, but did not reach statistical significance. However, this is interesting to consider along with the observation that mitochondrial P/O ratio is reduced in older sedentary adults. Both observations are in line with the “uncoupling to survive” theory, which posits that mitochondria become more uncoupled with age, and in this case sedentary aging, to prevent the deleterious effect of elevated ROS emission [53].

Taken together, the data clearly indicate that reduced physical activity, cardiovascular fitness and increased adiposity largely explain reductions in mitochondrial energetics in skeletal muscle from older adults [52]. Interestingly, VO_2_peak was lower in OA compared to YA, despite similar muscle mitochondrial energetics. This suggests a detrimental aging effect on other physiological determinants of VO_2_peak, potentially cardiac output or muscle perfusion [54], that is not completely abrogated by aerobic physical activity.

### Impact of chronological age and physical activity on intramyocellular lipid

Elevated intramyocellular lipid (IMLC) has been reported in vastus lateralis of older adults and can occur along with mitochondrial dysfunction and insulin resistance [55, 52]. We extend these findings to show that IMCL is elevated in soleus but not tibialis anterior muscle of OS compared to YA participants. It is not apparent why IMCL accumulation might occur specifically in soleus verses TA. However, soleus has a higher proportion of oxidative type I fibers [56] compared to the TA [57] and it has been shown that ectopic lipid accumulates to a greater extent in type I fibers of vastus lateralis compared to type II fibers in the context of obesity [58, 59], sedentary aging [52] and aging with metabolic syndrome [60]. In addition, type I fiber IMCL accumulation associates with poor contractile function [59]. Evidence in rodents confirm that soleus is particularly susceptible to the negative effects of physical inactivity (immobilization, hind limb unloading), on mitochondrial dysfunction, insulin resistance and loss of contractile function. Interestingly, some reports indicate that that the oxidative capacity of the TA is increased with aging, possibly to compensate changes in gait biomechanics [61, 62], which in turn could stave off age-associated IMCL accumulation. Taken together these reports suggest that soleus may be more susceptible to IMCL accumulation with sedentary behavior in aging and raises the interesting question that age and physical activity may have muscle group specific effects on IMCL content and subsequently on physical function.

### Impact of chronological age and physical activity on exercise efficiency and mitochondrial P/O ratio

Aging is also associated reduced exercise efficiency, defined as an increase in the energy expenditure of exercise, which likely contribute to impairments in physical activity [32, 31, 30]. Exercise efficiency can be improved with endurance training in both young [63] and older adults [32, 11]. Alterations in contractile coupling is the conventional explanation for differences in exercise efficiency [64], although a role for mitochondria in exercise efficiency has been previously suggested [65]. Others have shown that dietary nitrite improves mitochondrial and exercise efficiency during exercise [66, 67] and that markers of mitochondrial content and function [11], and the proportion of type I fibers [63] associate with exercise efficiency, all evidence suggesting that mitochondria play a role in mediation of exercise efficiency. In addition, our group previously reported that P/O ratio, calculated as ATP_Max_/state 3 respiration (PI+II), was related to 400m walk time in older adults [7]. Here, we extend those findings to show that mitochondrial P/O ratio is reduced in older sedentary adults when compared to young and old physically active groups, evidence indicating that reduced physical activity contributes to reduced mitochondrial efficiency in skeletal muscle from older adults. In addition, P/O ratio was related to both NE and GE during submaximal exercise, data which for the first time suggests that skeletal muscle mitochondrial efficiency is linked with whole body exercise efficiency in older adults. Alterations in mitochondrial efficiency of ATP production may be caused by altered inner mitochondrial membrane proton leak (uncoupling), possibly in response to greater ROS emission, or by reduced electron leak from the electron transport chain. Others have shown that mitochondrial uncoupling occurs in aging [68, 69], however further investigations into molecular mechanisms of mitochondrial uncoupling are needed to further understand the relationship with reduced exercise efficiency and likely the capacity for mobility in older adults.

### Mitochondrial energetics associate with muscle quality and physical function

Loss of muscle strength and quality underlie functional decline and predict mobility limitations in older adults [70, 71]. A recent report from Zane et al., showed that, in 326 participants from the Baltimore Longitudinal Study of Aging, mitochondrial phosphorylation capacity corresponds with muscle strength and quality in older adults [9]. Here, with a smaller cohort of older adults (n=29) but with a wide range of PA level, we further this observation to unveil that multiple aspects of mitochondrial energetics (ATPmax, P/O ratio, H_2_O_2_ emission) and muscle lipid content (IMAT and IMCL) strongly associate with muscle quality, supporting the hypothesis that they are likely important biological factors in loss of muscle quality with aging. In addition, multiple regression modelling showed that the combination of age, sex, ATPmax and IMCL together explained 60.6% of the variation of muscle quality in older adults.

These data are supported by preclinical evidence showing that mitochondrial reactive oxygen species can affect muscle mass by depressing protein synthesis [72-74] and impairing postprandial AKT phosphorylation, a major driver of the mTOR pathway and protein synthesis [44]. Genetic models of elevated mitochondrial ROS show a premature aging phenotype including loss of muscle mass and contractile properties [75]. Interestingly, soleus IMCL was significantly associated with many of the muscle and physical function measurements. In line with this observation, other have shown that intramyocellular lipids are linked with slower myofiber contraction, force and power development in obese older adults [59] and specific lipid species may mediate fatigue and weakness [76]. We also found that the combination of ATPmax, VO2peak and leg lean mass explained 54% of the variation in time taken to complete the repeated chair stand. These data support an emerging paradigm that cardiovascular and muscle bioenergetics play an important role in the etiology of mobility loss with aging [7-9], thus should be considered as a feasible therapeutic target to delay age-associated loss of muscle function and sarcopenia.

Our study has a few potential limitations that should be noted. We did not characterize all potential covariates that may influence the relationship between age and mitochondrial capacity and efficiency, including muscle fiber type. Also, while our study groups were balanced for sex ratio, we did not have enough statistical power to adequately investigate sex differences in aging, physical activity and mitochondrial energetics, likely an important biological factor. Thus, larger studies are needed to generate the next level of evidence that mitochondrial energetics are linked with physical function in men and women specifically. In addition, this study included few very low functioning people. Inclusion of more very low functioning older adults may have further strengthened the observed relationships between muscle mitochondrial capacity/efficiency and physical function. In summary, this study provides a number of novel and clinically relevant observations in well phenotyped young and older adults, including; 1) the variation in physical activity (predominantly endurance exercise), largely explains “age-associated” deficits in mitochondrial energetics and efficiency in older adults, 2) mitochondrial ATPmax and efficiency in skeletal muscle are linked with exercise efficiency, 3) skeletal muscle ATPmax nad IMCL are strongly associated with muscle contractile function and physical function in older adults.

## AUTHORS CONTRIBUTIONS

GD, RAS, XZ, EAC, HHC, and PMC contributed to study execution, researched the data and reviewed and approved the manuscript. FY and GD assisted with statistical analysis. GD contributed to data interpretation and writing the manuscript. All co-authors reviewed and approved the manuscript. PMC contributed to the study concept and design, statistical analysis, interpretation of the data and wrote the manuscript. PMC is the guarantor of the data.

## ACKNOWLEDGEMENTS

The authors gratefully appreciate the contributions of our study participants and acknowledge the excellent technical assistance of the imaging, recruitment, clinic, calorimetry, laboratory, and nutrition core staff at TRI-MD, Florida Hospital. Medical oversight for this study was provided by Richard E. Pratley, M.D. and Christian Meyer, M.D. This study was supported by funding from the National Institutes of Health | National Institute on Aging (K01 AG044437) awarded to PMC.

